# Differential gene expression profiling of lumbar disc degeneration in the RuDDS cohort

**DOI:** 10.64898/2026.11.23.768739

**Authors:** Aleksandr V. Tiapkin, Artemii A. Ivanov, Olga N. Leonova, Denis Maslov, Aleksandr V. Krutko, Alexey V. Peleganchuk, Iliya D. Isakov, Eugene A. Mushkachev, Tatiana S. Golubeva, Veniamin S. Fishman, Elizaveta E. Elgaeva, Yakov A. Tsepilov

## Abstract

Lumbar disc degeneration disease (LDDD) is a major structural cause of low back pain, one of the leading causes of disability worldwide, and has no disease-modifying treatment. We performed RNA sequencing of 72 intraoperative intervertebral disc samples from the Russian disc degeneration study (RuDDS) cohort, comparing severe (Pfirrmann grades IV–V) with moderate (grade III) degeneration. We identified 731 differentially expressed genes (DEGs; 347 upregulated and 384 downregulated in severely degenerated discs) in the 57 samples that passed quality control. Upregulated DEGs were enriched for extracellular matrix organisation, ossification, and TGF-*β* and Hippo signalling, whereas downregulated DEGs were enriched for immune and inflammatory processes. We identified five differentially active transcription factors, including the TGF-*β* effectors SMAD3 and SMAD4 and the osteogenic regulator DLX5. Of the five, only DLX5 was also a DEG. DEGs were enriched for LDDD-associated genes from Open Targets (odds ratio = 1.68, *P* = 3.37 *×* 10*^−^*^6^). Connectivity mapping against the LINCS L1000 database showed that histone deacetylase inhibitors and glucocorticoids reverse the signature of severe degeneration. Together, these results point to TGF-*β*-driven fibrotic and osteogenic remodelling as the dominant transcriptional feature of severe disc degeneration, and prioritise candidate genes and compounds for functional validation.

## Introduction

Low back pain is among the leading causes of disability worldwide ^1^, and lumbar disc degeneration disease (LDDD) is a major structural contributor to it ^2^. Treatment for LDDD remains symptomatic. TNF-*α* antagonists have been trialled for discogenic low back pain with limited success ^3^, and senolytics and NF-*κ*B inhibitors remain preclinical ^4–6^. Developing disease-modifying therapy requires molecular targets identified in degenerating human disc tissue.

The intervertebral disc (IVD) comprises a hydrated, proteoglycan-rich nucleus pulposus (NP) enclosed by the collagen lamellae of the annulus fibrosus (AF) ^7^. Degeneration depletes aggrecan and water, replaces type II collagen with type I, and converts the NP into a stiffer, fissured, fibrous tissue ^8^. Disc degeneration is graded on MRI using the Pfirrmann classification, from grade I (normal disc) to grade V (collapsed disc space) ^9^. Several processes are implicated in these changes, including extracellular matrix (ECM) breakdown, chronic inflammation, oxidative stress and cellular senescence ^10–13^. NP fibrosis is driven by chronic TGF-*β*/SMAD signalling ^14,15^, with further contributions from Hippo, NF-*κ*B and HIF-1*α* signalling ^16–19^, and NP cells can adopt an osteogenic phenotype ^20^. Single-cell studies have resolved the NP into chondrocyte-like, progenitor, stromal and immune subpopulations whose proportions shift with degeneration ^21–23^. Bulk RNA sequencing (RNA-seq) of disc tissue therefore conflates changes in cell composition with changes within individual cells.

Few studies have profiled the transcriptome of degenerating human lumbar disc tissue. A 2025 scoping review retained only 29 human tissue studies for synthesis, most of them small and cross-sectional ^19^. Jin et al. profiled 122 intraoperative NP samples by bulk RNA-seq and defined four molecular subtypes (C1, collagenesis; C2, ossification; C3, low chondrogenesis; C4, fibrogenesis). However, 94 of these samples were Pfirrmann grades IV–V, leaving only two grade II and 26 grade III discs to represent milder degeneration ^24^.

Here we performed a differential expression analysis between moderate (Pfirrmann grade III) and severe (grades IV–V) degeneration in intraoperative IVD samples from the multi-centre Russian disc degeneration study (RuDDS) cohort ^25^. We characterised sample composition by deconvolution against a healthy IVD single-cell atlas and by molecular subtyping, and inferred transcription factor (TF) and microRNA (miRNA) activity. Differentially expressed genes (DEGs) were compared with Open Targets and genome-wide association study (GWAS) evidence, and Library of Integrated Network-Based Cellular Signatures (LINCS) L1000 connectivity data were queried for compounds that reverse the LDDD signature. These analyses extend beyond a list of individual DEGs to the regulators that organise the degenerative transcriptome, and prioritise candidates for functional validation and drug development.

## Results

### Cohort description

The RuDDS cohort is a multi-omics biobank of patients with lumbar disc degeneration ^25^. It recruits adult patients with lumbar spine MRI at two clinical centres, in Novosibirsk and Moscow, and collects clinical, socio-demographic and MRI data, whole blood and plasma from all participants, and intervertebral disc tissue from those who undergo spinal surgery. All intraoperative disc samples analysed in this study were collected at the Novosibirsk centre. Disc degeneration is common even in asymptomatic adults, with an MRI prevalence rising from 37% at age 20 to 96% at age 80^26^, so healthy discs are rarely available for comparison. The cohort therefore defines controls conditionally, as discs with less severe degeneration.

Before quality control, RNA-seq data were available for 72 samples (40 cases and 32 conditional controls). The epidemiological characteristics of the samples are presented in Table 1. Cases included a slightly higher proportion of patients assigned male at birth. Age and RNA integrity number (RIN) did not differ statistically significantly between cases and controls before or after quality control. Body mass index (BMI) did not differ before quality control, but after quality control cases had a higher mean BMI than controls (29.0 vs. 25.8 kg/m^2^; Welch’s *t* -test, *P* = 0.01).

**Table 1.** RuDDS cohort summary before and after quality control.

|  | **Before QC** |  | **After QC** |  |
|---|---|---|---|---|
|  | **Control** | **Case** | **Control** | **Case** |
| Total samples | 32 | 40 | 24 | 33 |
| % Male | 37.50 | 50.00 | 45.83 | 48.48 |
| Mean age ± SD (years) | 45.88 ± 11.85 | 45.23 ± 13.85 | 45.38 ± 11.42 | 45.45 ± 14.62 |
| Mean BMI ± SD (kg/m<sup>2</sup>) | 26.83 ± 4.19 | 29.02 ± 5.61 | 25.75 ± 4.06 | 29.02 ± 5.20 |
| Mean RIN ± SD | 7.86 ± 0.87 | 7.85 ± 0.98 | 7.95 ± 0.92 | 8.04 ± 0.96 |

QC, quality control; RIN, RNA integrity number; BMI, body mass index; SD, standard deviation.

### RNA extraction and quality control

We extracted high-quality RNA from 72 samples (mean RIN = 7.9). The raw sequencing data quality was adequate, with a mean sequencing depth of 23.9 million read pairs and mean Phred quality scores of 35.8 (read 1) and 35.5 (read 2). On average, 99.1% of reads were mapped to the genome (*Supplementary Table 1*). Of all reads, 8.7% did not overlap any annotated feature and 3.3% were ambiguous, so 87.1% were assigned to gene annotation features (exons).

To identify potential outliers in the RNA-seq data, we performed a principal component analysis (PCA; Materials and Methods). We identified 15 outliers (7 case samples and 8 conditional controls; *Supplementary Fig. 1*; *Supplementary Note 1*). To account for unmeasured sources of variation, such as differences in tissue composition and sample handling, we included two surrogate variables (SV1 and SV2) in the DESeq2 design (Materials and Methods; *Supplementary Fig. 1E*).

**Figure 1.**
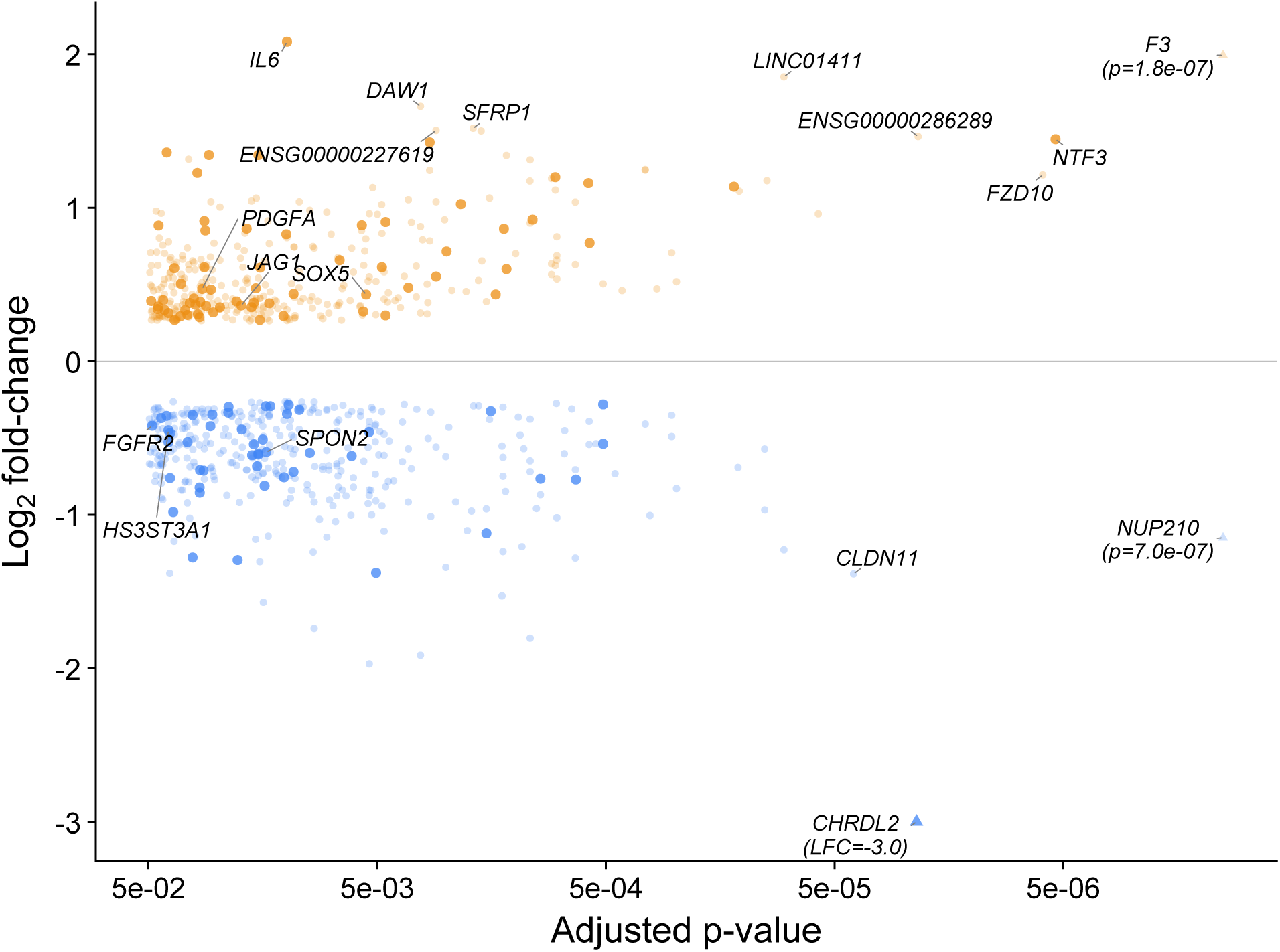
Volcano plot of differentially expressed genes. Log2 fold-change is plotted against adjusted *P*-value (reversed log_10_ scale) for the 731 DEGs. Orange and blue points indicate upregulated and downregulated DEGs, respectively; larger, more opaque points mark Open Targets LDDD-associated genes. Genes are labelled if they meet at least one of three criteria: an Open Targets score *≥* 0.1; log_2_ FC *>* 1.5 or *< −*2; or adjusted *P <* 5 *×* 10*^−^*^5^. The *x* -axis is capped at 10*^−^*^6^ and the *y* -axis at log_2_ FC = *−*3; genes beyond either cap (*F3*, *NUP210* and *CHRDL2*) are shown as triangles annotated with their exact adjusted *P*-value or log_2_ fold-change.

### Cell-type composition and sample heterogeneity

To characterise the cellular composition of RuDDS IVD samples and to contextualise the observed heterogeneity, we performed bulk RNA-seq deconvolution using BayesPrism ^27^ with the Gan et al. single-cell IVD atlas of healthy discs ^21^ as the reference (*Supplementary Fig. 2*; *Supplementary Table 2*). The atlas comprised nine cell types: three chondrocyte subpopulations (Chond1, Chond2, Chond3), notochordal cells (Noto), nucleus pulposus progenitor cells (NPPC), Stroma, endothelial cells (EC), Pericyte, and Blood.

Across all 72 samples, Stroma and Chond2 were the two most abundant inferred cell types. All remaining populations were present at substantially lower median proportions (*Supplementary Fig. 2*). Several samples identified as outliers during PCA-based quality control displayed elevated Blood or Pericyte proportions, consistent with contamination of the surgical specimen with peripheral blood or perivascular tissue.

Hierarchical clustering of the cell-type proportions partitioned the 72 samples into three groups (*Supplementary Fig. 2*): a Chondrocyte-rich group (D1, *n* = 33) with high Chond2 proportions, a Stroma-rich group (D2, *n* = 33) and a Blood-rich group (D3, *n* = 6) consisting entirely of outliers identified by PCA-based quality control (*Supplementary Fig. 1*; *Supplementary Note 1*).

**Figure 2.**
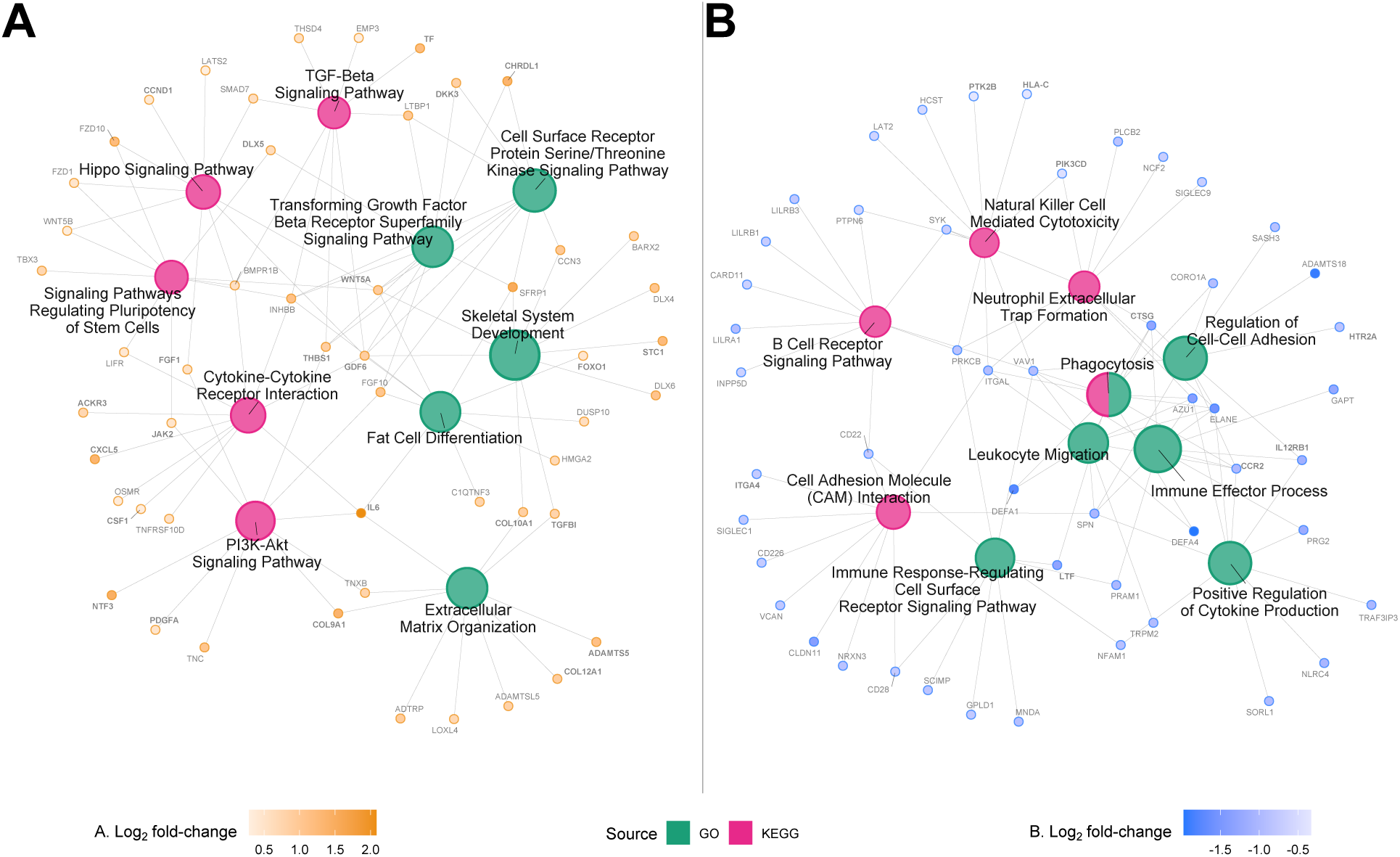
Over-representation analysis of upregulated and downregulated DEGs. The five GO Biological Process categories and five KEGG pathways with the lowest adjusted *P*-values, and their member genes, for (A) upregulated and (B) downregulated DEGs. Redundant GO categories were removed before selection. Category node size increases with the number of member DEGs, and node colour indicates the source database. Each category shows up to ten member DEGs with the largest absolute log_2_ fold-change. Gene nodes are coloured by log_2_ fold-change, and gene labels in bold mark Open Targets LDDD-associated genes.

We assigned samples to the molecular subtypes of Jin et al. ^24^ by hierarchical clustering on their 248-gene subtype classifier panel (Materials and Methods; *Supplementary Figs 3 and 4*). The four clusters of the full cohort corresponded to subtypes C4 (*n* = 26), C1 (*n* = 25), C2 (*n* = 16) and C3 (*n* = 5). In Jin et al., these subtypes are characterised by inflammatory fibrosis (C4), chondrogenic matrix synthesis (C1), osteogenic remodelling (C2) and mitochondrial respiratory metabolism with reduced chondrogenesis (C3). All C3 samples were outliers from the Blood-rich group and showed coordinated high expression of haemoglobin genes. All C1 samples were Chondrocyte-rich and 25 of 26 C4 samples Stroma-rich, whereas C2 samples were split between the two (7 and 8), with the remaining C2 sample, an outlier, in the Blood-rich group (adjusted Rand index between subtypes and deconvolution groups, 0.57). Among the 57 samples passing quality control, clustering yielded subtypes C1 (*n* = 23), C4 (*n* = 19) and C2 (*n* = 15), with the same correspondence to deconvolution groups (*Supplementary Fig. 5A*). C2 included a higher proportion of case samples (13 of 15) than C1 (10 of 23). The two surrogate variables of the DESeq2 model captured most of this heterogeneity (*Supplementary Fig. 5B, C*). Jointly, SV1 and SV2 explained 64% of the variance in deconvolution group and 87% and 71% in Chond2 and Stroma proportions, and were strongly associated with subtype.

### Differentially expressed genes and pathway enrichment

We obtained the differential gene expression profile for 15,144 protein-coding or long non-coding RNA genes (*Supplementary Table 3*). We identified 731 DEGs (347 upregulated and 384 downregulated in severely degenerated discs) with absolute fold-change greater than 1.2 (*|* log_2_ FC*| >* log_2_(1.2) *≈* 0.26) and adjusted *P <* 0.05, comprising 658 protein-coding genes and 73 long non-coding RNA genes (Fig. 1; *Supplementary Table 4*).

Out of 1,865 genes associated with LDDD extracted from the Open Targets Platform ^28^, 1,510 were in the background gene list. Fisher’s exact test indicated statistically significant enrichment of LDDD-associated genes in our DEG list (odds ratio = 1.68, *P* = 3.37 *×* 10*^−^*^6^) with an overlap of 112 genes (*Supplementary Table 4*). Six overlapping genes were strongly associated with LDDD (Open Targets score *≥* 0.1): *HS3ST3A1*, *JAG1*, *PDGFA*, *FGFR2*, *SOX5*, *SPON2*.

Over-representation analysis of Gene Ontology (GO) Biological Process categories and Kyoto Encyclopedia of Genes and Genomes (KEGG) pathways identified 57 enriched categories among upregulated and 50 among downregulated DEGs (*Supplementary Tables 5 and 6*). Upregulated DEGs were enriched for skeletal system development and ECM organisation, together with the TGF-*β*, Hippo and PI3K–Akt signalling pathways (Fig. 2A), as well as ossification and osteoblast differentiation (*Supplementary Table 5*). Downregulated DEGs were enriched for immune processes, including immune effector process, phagocytosis and B cell receptor signalling (Fig. 2B), as well as regulation of inflammatory response (*Supplementary Table 6*).

Gene set enrichment analysis (GSEA) identified 22 KEGG pathways with statistically significant enrichment (*Supplementary Table 7*). Hippo and TGF-*β* signalling showed the strongest positive enrichment, and lysosome biogenesis and neutrophil extracellular trap formation the strongest negative enrichment.

### Transcription factor activity inference

Using decoupleR ^29^, we identified five TFs exhibiting statistically significant differential activity (Fig. 3; *Supplementary Table 8*). This activity is inferred by comparing the observed expression changes across the full differential gene expression profile against prior knowledge of TF-target interactions from the CollecTRI network ^30^. A TF is considered ‘activated’ when its targets show a collective expression pattern consistent with that TF’s predicted influence, whereas an ‘inactivated’ TF shows a pattern opposed to its predicted influence.

**Figure 3.**
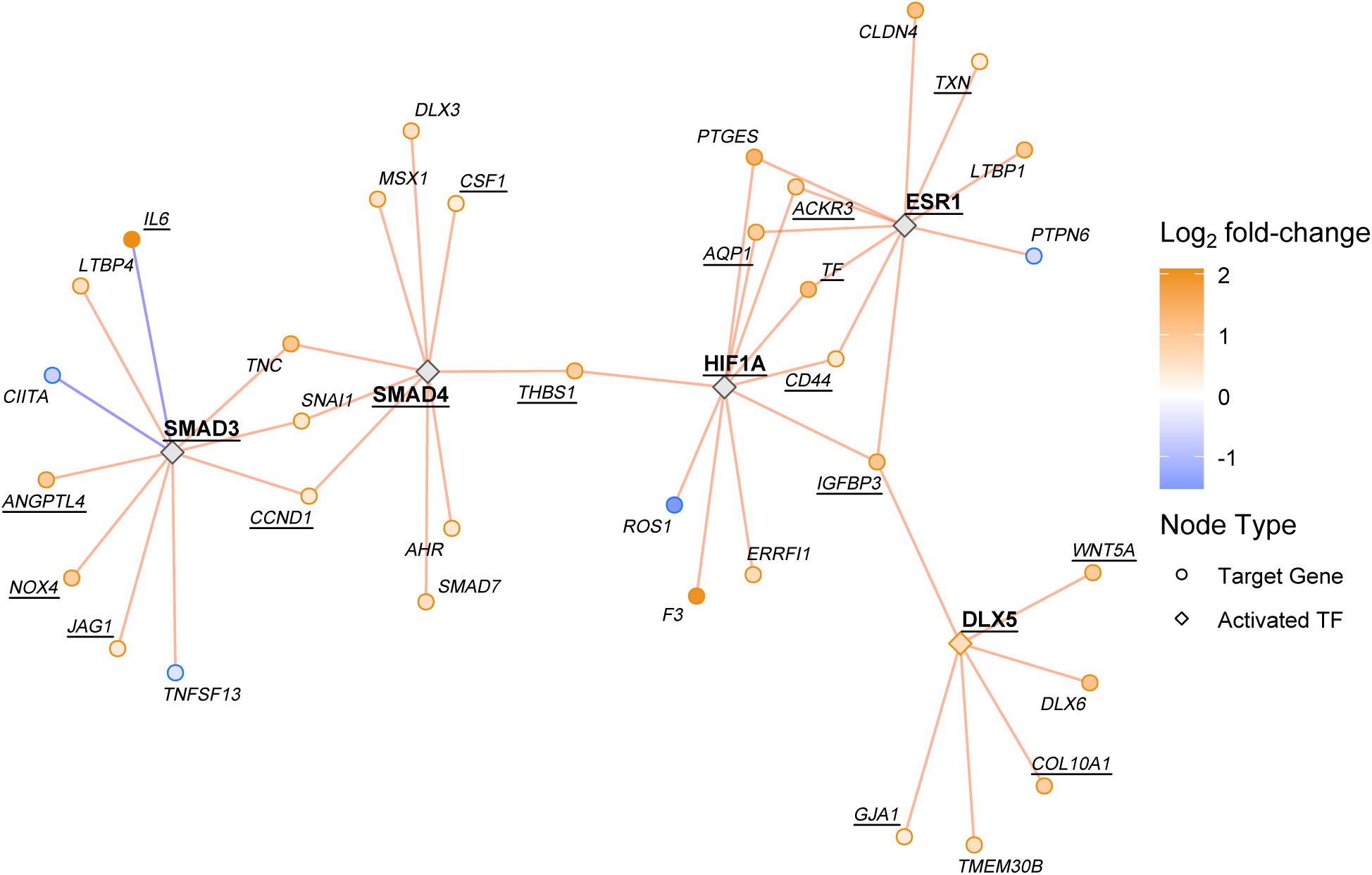
Regulatory network of differentially active transcription factors and their top target genes. The five activated TFs (diamonds) and their ten most strongly contributing target DEGs (circles) from the CollecTRI prior knowledge network ^30^. Target nodes are coloured by log_2_ fold-change. Orange edges indicate activation and blue edges repression. Underlined labels mark Open Targets LDDD-associated genes.

All five differentially active TFs (DLX5, ESR1, HIF1A, SMAD3, SMAD4) were activated, and each had a prior association with LDDD in Open Targets. Only *DLX5* was itself differentially expressed (log_2_ FC = 0.61), although every TF gene was expressed in our samples, with a mean transcripts per million across samples ranging from 1.5 to 189.7. Based on prior knowledge, these TFs are predicted to regulate 66 DEGs (51 upregulated, 15 downregulated) with 94 TF-target interactions.

Among the DEG targets of activated TFs we found genes with the largest absolute fold-changes (| log_2_ FC*| >* 1), such as *IL6*, *F3*, *ROS1*, *PTGES* and *TNC*.

### miRNA target enrichment analysis

GSEA against miRDB v8 predicted miRNA target gene sets ^31^ from the Molecular Signatures Database (MSigDB) C3 MIR:MIRDB collection ^32^ identified 197 target sets enriched at a false discovery rate (FDR) below 1% (*Supplementary Table 9*). Of these, 194 had a positive normalised enrichment score (NES), placing the targets of these miRNAs among upregulated genes. Three (miR-4505, miR-5787 and miR-6825-5p) had a negative NES.

Targets of miR-338-5p had the highest NES (2.10), with a leading edge that included *IL6*, *CCN2* and *NTF3*. The leading edges of the miR-29 family (miR-29a-3p, NES = 1.55; miR-29b-3p/miR-29c-3p, NES = 1.53) were rich in ECM genes, including *COL1A1*, *COL1A2*, *COL3A1*, *COL5A1*, *COL11A1*, *LOX*, *LOXL2* and *FBN1*, and miR-5682 (NES = 1.84) showed a similar collagen-rich leading edge. Enriched sets also included targets of the miR-548 family (NES = 1.46–2.10; *TNC*, *RUNX2*, *TGFBR1*), miR-624-5p (NES = 1.92; *F3*, *TGFBR1*) and miR-145-5p (NES = 1.58; *ADAMTS5*, *TGFB2*, *SOX11*).

### Connectivity mapping against LINCS L1000

To generate hypotheses about compounds that may reverse the LDDD transcriptional programme, we queried our DEG signature against the LINCS L1000 perturbagen database ^33^ using the gess_lincs function of the *signatureSearch* Bioconductor package ^34^ (*Supplementary Table 10*). The query comprised the 233 upregulated and 246 downregulated DEGs present in the LINCS reference database. A negative normalised connectivity score (NCS) indicates that a compound’s transcriptional effect is anti-correlated with the LDDD signature, making it a reversal candidate. A positive NCS marks a mimic. At a weighted connectivity score (WTCS) FDR below 1%, we identified 419 compound–cell-line entries, comprising 305 reversal candidates and 114 mimics.

The top-ranked reversal candidate was rilmenidine (NCS = *−*1.78), an *α*_2_-adrenergic and imidazoline receptor agonist. Histone deacetylase (HDAC) inhibitors were the most frequent drug class, with 12 of 13 entries reversing the signature across four cell lines (trichostatin A, panobinostat, vorinostat, scriptaid, HC-toxin, apicidin). All six glucocorticoid entries reversed the signature (NCS = *−*1.32 to *−*1.51), including betamethasone and triamcinolone. Other recurring reversal classes included EGFR inhibitors (afatinib, lapatinib, erlotinib, neratinib), SRC kinase inhibitors (saracatinib, WH-4023, PP-2), dopamine receptor antagonists (perphenazine, trifluoperazine, ziprasidone) and adrenergic modulators. The p38 MAPK inhibitor SB-202190 (NCS = *−*1.46) and the ROCK inhibitor GSK-429286A (NCS = *−*1.49) each reversed the signature in two cell lines. Reversal candidates further included the natural compounds epicatechin gallate (NCS = *−*1.68), *α*-linolenic acid (NCS = *−*1.58) and apigenin (NCS = *−*1.55).

Mimic candidates included the cardiac glycosides helveticoside (NCS = +1.67), the top-ranked mimic, and gitoxigenin (NCS = +1.50), as well as icosapent (eicosapentaenoic acid; NCS = +1.45) and the vitamin D analogue calcipotriol (NCS = +1.41).

## Discussion

We compared severe (Pfirrmann grades IV–V) with moderate (grade III) degeneration in intraoperative disc tissue from the RuDDS cohort, complementing the larger subtype analysis of Jin et al. ^24^, which comprised mostly cases (grades IV–V). To our knowledge, with 72 sequenced discs, this is among the largest case-control RNA-seq studies of intraoperative disc material.

Intraoperative disc material was highly heterogeneous in cell composition. To control for this heterogeneity, we included two surrogate variables in the differential expression model. They absorbed most of the compositional variation, including the proportions of stromal and chondrocytic material (*Supplementary Fig. 5*). Although NP and AF were not separated before sequencing, deconvolution indicated predominantly NP material. Chond1, which localises mainly to AF and cartilage endplate in the reference atlas ^21^, was nearly absent from our samples (mean 0.6%).

The compositional groups corresponded to the transcriptomic subtypes of Jin et al. ^24^. Chondrocyterich samples matched the C1 subtype, which is characterised by type II collagen and aggrecan synthesis. Stroma-rich samples matched the fibrogenic, inflammatory C4 subtype, in which collagens, particularly type I collagen, predominate over proteoglycans in the matrix ^24^. Samples with a Blood signature were most closely related to the C3 subtype, with high expression of *S100A8* and *S100A9*, which are abundant in neutrophils and monocytes (*Supplementary Fig. 3*). This signature likely reflects peripheral blood retained in the surgical specimen, and we excluded these samples from the differential expression analysis. Cases dominated the osteogenic C2 subtype, the only subtype split evenly between Chondrocyte-rich and Stromarich samples.

TGF-*β* signalling was the most consistent signal across our analyses. It was among the most strongly enriched pathways, and two of the activated TFs, SMAD3 and SMAD4, transduce TGF-*β* signalling and induce type I collagen expression ^35^. Excessive TGF-*β* activation promotes disc degeneration ^36^. In NP cells, TGF-*β*1 signalling through SMAD2/3–SMAD4^37^ induces a myofibroblast-like phenotype that replaces the proteoglycan-rich matrix with type I and III collagen ^15^. Consistent with TGF-*β* activity, targets of miR-145-5p, a miRNA that TGF-*β* suppresses ^38^ and that protects NP cells from apoptosis and matrix degradation ^39,40^, were upregulated.

DLX5 and HIF1A, the two other activated TFs, point to osteogenic and hypoxic responses. DLX5, the only TF whose gene was itself differentially expressed, is an osteogenic master regulator ^41^. Its expression in human NP tissue increases with Pfirrmann grade, and in cultured NP cells DLX5 inhibited SMAD2/3 phosphorylation ^42^. In our tissue samples, by contrast, DLX5, SMAD3 and SMAD4 all showed increased inferred activity. *DLX5* is also among the most tissue- and cell-type-specific genes in Open Targets baseline expression data, whereas *SMAD3*, *SMAD4* and *HIF1A* are expressed ubiquitously ^28^. HIF1A maintains NP cell survival under hypoxia and limits oxidative stress ^12,43^, so its activation may reflect a compensatory response to inflammatory stress. Inflammation persisted in severe degeneration, although the higher expression of immune-cell genes in grade III discs suggests greater leukocyte infiltration at the moderate stage. *IL6* and *PTGES*, both targets of the activated TFs, were among the most strongly upregulated DEGs, consistent with cytokine and prostaglandin production by disc cells themselves ^11^.

All six DEGs with the strongest Open Targets association with LDDD (*HS3ST3A1*, *JAG1*, *PDGFA*, *FGFR2*, *SOX5*, *SPON2*) have GWAS evidence for disc degeneration or herniation ^28^. *SOX5* lies at one of the strongest GWAS loci for intervertebral disc degeneration ^44^ and back pain ^45^. Beyond these six, *IGFBP3*, located near a GWAS signal for surgically treated disc herniation ^44^, was also upregulated in an independent AF transcriptome ^46^. Drug mechanisms with genetic support are 2.6 times more likely to reach approval ^47^, so genetic evidence adds independent support for these genes as therapeutic targets.

Connectivity mapping nominated several compounds with established effects on disc cells or on patients with disc herniation. Glucocorticoids formed a consistent reversal class that included betamethasone and triamcinolone, both used in epidural injections for radicular pain from disc herniation ^48^. Two further candidates protect the disc in experimental models. The p38 MAPK inhibitor SB-202190 reversed the signature, and p38 MAPK inhibitors reduce inflammatory mediator production and matrix degradation in NP cells ^49^. Apigenin attenuates puncture-induced disc degeneration in rats by restoring autophagic flux ^50^. Rilmenidine, an imidazoline receptor agonist and the top-ranked candidate, and HDAC inhibitors, the most frequent reversal class, gave the strongest reversal signals and are priority candidates for testing in NP cells.

This study has several limitations. Control samples had grade III degeneration, not healthy discs, so the analysis cannot detect expression changes at earlier stages. The DEGs therefore reflect progression from moderate to severe degeneration, not disease onset. The samples are highly heterogeneous in cell composition, and bulk RNA-seq cannot assign expression changes to specific cell types. Single-cell or spatial profiling of intraoperative tissue is needed to increase the resolution of the analysis. Poly-A-selected libraries do not capture miRNAs, so miRNA activity was inferred from the expression of predicted targets. L1000 profiles come from cancer and other non-IVD cell lines, so connectivity scores may not reproduce drug responses in disc cells. The candidate regulators and compounds require functional testing in patient-derived NP cells.

In conclusion, severely degenerated discs showed upregulation of genes for fibrotic and osteogenic ECM remodelling, whereas immune-cell genes were more highly expressed in grade III discs. Pathway and regulator analyses pointed to TGF-*β* signalling as a likely driver of severe degeneration. This study provides a transcriptomic resource for intraoperative LDDD research and a set of candidate regulators and compounds for future mechanistic and drug development studies.

## Materials and Methods

### Study design and sample collection

Following the previously published protocol ^25^, IVD fragments were obtained from patients who underwent spinal surgery at the Novosibirsk Research Institute of Traumatology and Orthopedics (Novosibirsk, Russia). The intraoperative material was placed in sterile 50 ml tubes and frozen in liquid nitrogen within 45 minutes of disc removal. It was then transported to the Institute of Cytology and Genetics (Novosibirsk, Russia) in liquid nitrogen and stored at *−*80 *^◦^*C.

### RNA extraction and sequencing

Because high-quality RNA is difficult to extract from IVDs ^51–53^, we used the extraction protocol of Ivanov et al. ^54^. Intraoperative IVD fragments were available from 72 samples, including 32 conditional control samples (Pfirrmann grade III) and 40 case samples (Pfirrmann grades IV–V). Strand-specific paired-end RNA-sequencing was performed on the DNBSEQ-G400 platform with a read length of 2 *×* 100 bp.

### Raw sequencing data processing and quality control

The analysis was implemented as a Nextflow ^55^ v26.10.5 pipeline (see Code Availability). The quality of raw sequencing data was assessed using FastQC ^56^ v0.12.1 and RNA-SeQC ^57^ v2.3.5 with the legacy counting mode (flag --legacy). Alignment and all subsequent analyses were carried out with R v4.5.1. Reads were aligned to the human reference genome (version GRCh38.p14) with GENCODE annotation v47 using the align function from Rsubread ^58^ v2.22.1 with default parameters. Gene-level counts were obtained with the Rsubread featureCounts function using default parameters and reversely stranded read counting (parameter strandSpecific = 2).

### Cell-type deconvolution

Cell-type composition of bulk RNA-seq samples was estimated using the BayesPrism ^27^ R package v2.2.2. As a single-cell reference we used the healthy human IVD atlas published by Gan et al. ^21^ (Gene Expression Omnibus (GEO) accession GSE160756), covering NP, cartilage endplate and AF compartments from five donors. Cell-type labels from the published cluster annotations defined nine populations: Chond1, Chond2, Chond3, Noto, NPPC, Stroma, EC, Pericyte and Blood. Prior to deconvolution, both matrices were restricted to protein-coding genes shared between the bulk and single-cell data. Ribosomal, mitochondrial, haemoglobin, sex-chromosome, actin, and *MALAT1* transcripts were removed. To identify compositionally distinct sample groups, the proportion matrix was hierarchically clustered by Ward’s method (ward.D2) on Euclidean distances, with the dendrogram cut at *k* = 3 for all 72 samples and at *k* = 2 for the 57 samples passing quality control.

### Molecular subtype classification

Transcriptomic heterogeneity of RuDDS samples was assessed relative to the four-subtype LDDD molecular classification of Jin et al. ^24^. Counts were transformed with the DESeq2 variance-stabilising transformation (blind = TRUE) for all 72 samples and for the 57 samples passing quality control. Samples were clustered on the 248-gene subtype classifier panel of Jin et al. (their Supplementary Table 3), of which 244 genes were present in the RuDDS data. Each gene was z-scored across samples, and samples were clustered hierarchically by Ward’s method (ward.D2) using Pearson correlation distance. The dendrogram was cut at *k* = 4 for all 72 samples and at *k* = 3 for the 57 samples passing quality control. Each cluster was assigned the Jin subtype whose marker genes (Fig. 2e of Jin et al.) and classifier genes it expressed most highly.

We tested whether the two surrogate variables of the DESeq2 model captured this heterogeneity by relating SV1 and SV2 in the 57 samples to deconvolution group (*k* = 2), estimated subtype, condition, and the estimated proportions of Chond2, Stroma, NPPC, EC and Blood cells. Pairwise associations were measured by Pearson correlation, with condition, deconvolution group and each subtype coded as binary indicators. The variance each variable shares with a single SV was estimated as the *R*^2^ of a linear regression of the SV on that variable and tested with an *F* test. The variance shared with both SVs jointly was estimated as the squared first canonical correlation between the variable and SV1 and SV2, and tested with a multivariate analysis of variance (Pillai’s trace). All *P*-values for these associations are nominal.

### Differential gene expression analysis

We used the DESeq2^59^ R package v1.48.2 to perform differential expression analysis. We filtered out low-count genes and retained only those with at least 25 counts in at least 15 samples.

To identify outliers among the samples, we applied the variance-stabilising transformation with blind dispersion estimation to the count matrix and performed PCA using the DESeq2::plotPCA function. Based on manual inspection of the resulting plots, 15 samples out of 72 were marked as outliers and removed from the analysis (*Supplementary Note 1*).

The initial DESeq2 design formula included *Sex* and *Age* as covariates and *Condition* (case-control status) as the variable of interest.

Extreme individual count outliers were identified using Cook’s distances provided by DESeq2 (0.02% of all values) and downweighted, reducing their impact on the analysis (*Supplementary Note 2*).

Hidden batch effects were modelled with surrogate variables estimated by the svaseq function from the sva v3.56.0 R package ^60,61^. To preserve the effect of *Condition* while retaining effects of known covariates, we included *Condition*, *Sex* and *Age* variables in the full model matrix and only *Sex* and *Age* variables in the reduced null model matrix when calculating surrogate variables. The optimal number of surrogate variables was determined using the num.sv function with Leek’s method. To approximate the impact of adding these surrogate variables to the DESeq2 design formula, we adjusted the variance-stabilised matrix to SV1 and SV2 using the removeBatchEffect function from the limma v3.64.3 R package ^62^ and visualised the results using PCA plots (*Supplementary Fig. 1E*). The adjusted matrix was used for visualisation purposes only.

Finally, the *DESeq* function was called with the following design formula:

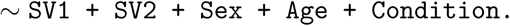

*P*-values were adjusted using the Benjamini–Hochberg procedure for multiple testing correction ^63^. Genes with an adjusted *P*-value below 0.05 and an absolute fold-change exceeding 1.2 (*|* log_2_ FC*| >* log_2_(1.2) *≈* 0.26) were considered differentially expressed. Subsequently, we restricted the analysis to protein-coding and long non-coding RNA genes.

### Gene ontology and pathway enrichment analysis

We performed over-representation analysis ^64^ separately on the upregulated and downregulated DEGs. GO Biological Process categories ^65,66^ were tested with enrichGO and KEGG pathways ^67^ with enrichKEGG, both from the clusterProfiler ^68^ R package v4.16.0. The full differential gene expression profile of 15,144 protein-coding and long non-coding RNA genes (*Supplementary Table 3*) served as the background. Gene sets of at least 20 genes were tested, and categories with a Benjamini–Hochberg adjusted *P*-value below 0.05^63^ were considered enriched. We removed redundant GO categories with the clusterProfiler simplify function at a semantic similarity cutoff of 0.7. For Fig. 2, the redundancy of GO categories was further reduced with a similarity cutoff of 0.35, and the five GO categories and five KEGG pathways with the lowest adjusted *P*-values were plotted with the enrichplot ^69^ R package v1.28.4.

### Functional class scoring

GSEA ^70^ of KEGG pathways was run on the full differential gene expression profile ranked by the DE-Seq2 Wald statistic, using clusterProfiler gseKEGG (DOSE implementation ^71^, 10,000 permutations) and a Benjamini–Hochberg adjusted *P*-value threshold of 0.01.

### Open Targets Platform data

LDDD-associated genes were retrieved from the publicly available Open Targets Platform ^28^ version 26.03 for three disease terms: lumbar disc degeneration (EFO_0004994), intervertebral disc degeneration (HP_0008419) and lumbar disc herniation (EFO_1002005). We took the union of the three gene lists, retaining the highest association score for genes present in more than one list. Enrichment of these genes among the DEGs was tested with Fisher’s exact test against the full differential gene expression profile.

### Transcription factor activity inference

Differential TF activity was inferred using the decoupleR ^29^ package v2.14.0 with the signed CollecTRI prior knowledge network ^30^ and the univariate linear model method (run_ulm), using the DESeq2 Wald statistic of the full differential gene expression profile as input. TFs with at least 15 targets in the profile were tested. A TF was considered differentially active if its Benjamini–Hochberg adjusted *P*-value was below 0.01 and its DEG targets changed in the direction predicted by its inferred activity. A DEG target was counted as consistent when activated targets of an activated TF were upregulated and repressed targets downregulated, or the reverse for an inactivated TF. We required at least five consistent DEG targets, a consistent fraction above 0.75 and a one-sided binomial *P*-value below 0.05 for that fraction against 0.5. The network in Fig. 3 was drawn with igraph v2.1.4 and ggraph v2.2.2 and shows, for each differentially active TF, the 10 DEG targets with the largest absolute Wald statistic.

### miRNA target enrichment analysis

To identify miRNAs whose predicted targets are enriched among up- or downregulated genes, we performed GSEA against the MSigDB ^32^ C3 MIR:MIRDB gene set collection (miRDB v8 predicted targets ^31^) retrieved using the msigdbr ^72^ R package v26.1.0. GSEA was performed with the fgsea^73^ R package v1.34.2 on the full differential gene expression profile ranked by the DESeq2 Wald statistic. Gene sets with a Benjamini–Hochberg adjusted *P*-value below 0.01 were considered statistically significant.

### Connectivity mapping

To identify compounds predicted to reverse the LDDD transcriptional signature, we queried the LINCS L1000 Phase I database ^33^ (GEO GSE92742; ExperimentHub identifier EH3226) using the gess_lincs function of the *signatureSearch* ^34^ package v1.22.0. The query signature comprised all upregulated and downregulated DEGs present among the 12,328 genes of the reference database. Compound–cell-line signatures with a WTCS FDR below 0.01 were retained and ranked by NCS. Compounds with a negative NCS induce an opposing gene expression profile and were classified as reversal candidates, whereas a positive NCS indicates a disease-mimicking profile.

### Ethical approval

The study was conducted in accordance with the Declaration of Helsinki. The study protocol was approved by the Local Ethical Committee of the Novosibirsk Research Institute of Traumatology and Orthopedics (No. 034/20, dated 2 October 2020) and by the Local Ethical Committee of the Priorov Central Research Institute of Traumatology and Orthopedics (CITO; Moscow, Russia; No. 1/21, dated 25 February 2021). All participants provided written informed consent.

## Author Contributions

A.V.T., O.N.L., E.E.E. and Y.A.T. conceived the study. A.V.T., A.A.I., D.M., T.S.G., E.E.E. and Y.A.T. developed the methodology. A.V.T. and Y.A.T. wrote the software, performed the formal analysis and wrote the original draft. A.A.I., O.N.L., A.V.K., A.V.P., I.D.I., E.A.M. and V.S.F. contributed to sample collection and sequencing. O.N.L., A.V.K., A.V.P., I.D.I. and E.A.M. provided patients, surgical material and clinical data. A.V.T., E.E.E. and Y.A.T. curated the data. A.V.T. prepared the figures. Y.A.T. supervised the study. E.E.E. and Y.A.T. administered the project. O.N.L., E.E.E. and Y.A.T. acquired funding. All authors reviewed and edited the manuscript and approved the final version.

## Data Availability

The raw RNA-seq data generated in this study are not publicly available and cannot be deposited in a public repository because of restrictions on sharing human genetic data. The full differential gene expression results are provided in *Supplementary Table 3*.

## Code Availability

The analysis code is available on GitHub at https://github.com/al-t1/rudds-rnaseq.

