## Supplementary Notes for "Differential gene expression profiling of lumbar disc degeneration in the RuDDS cohort"

### Supplementary Information

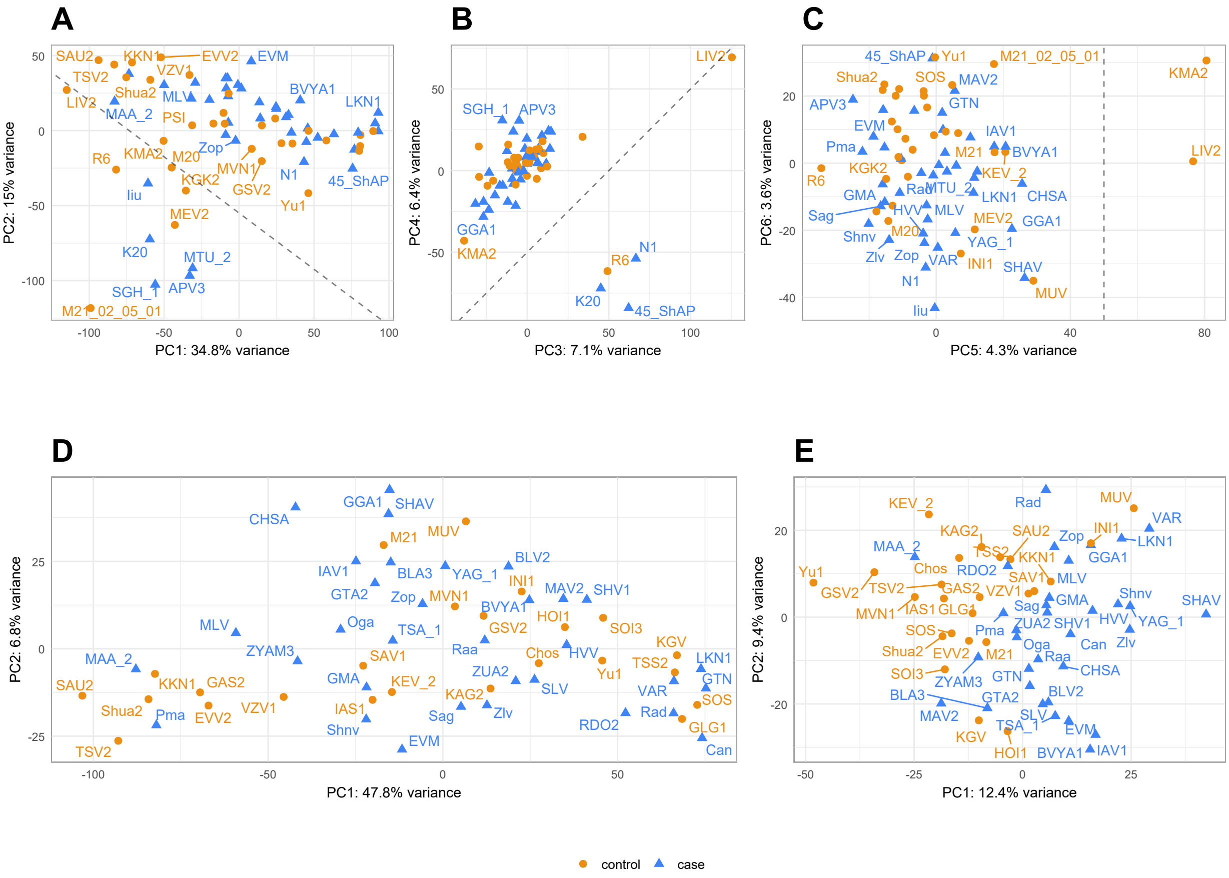


**Supplementary Fig. 1 | Sample quality control.** (A–C) Principal component analysis (PCA) plots of expression profiles across all samples for principal components (A) 1 and 2, (B) 3 and 4, and (C) 5 and 6. (D) PCA plot of components 1 and 2 after outlier filtering. (E) PCA plot of components 1 and 2 after outlier filtering and adjustment for the first two surrogate variables using the removeBatchEffect function from the limma R package. Dashed lines separate outliers.

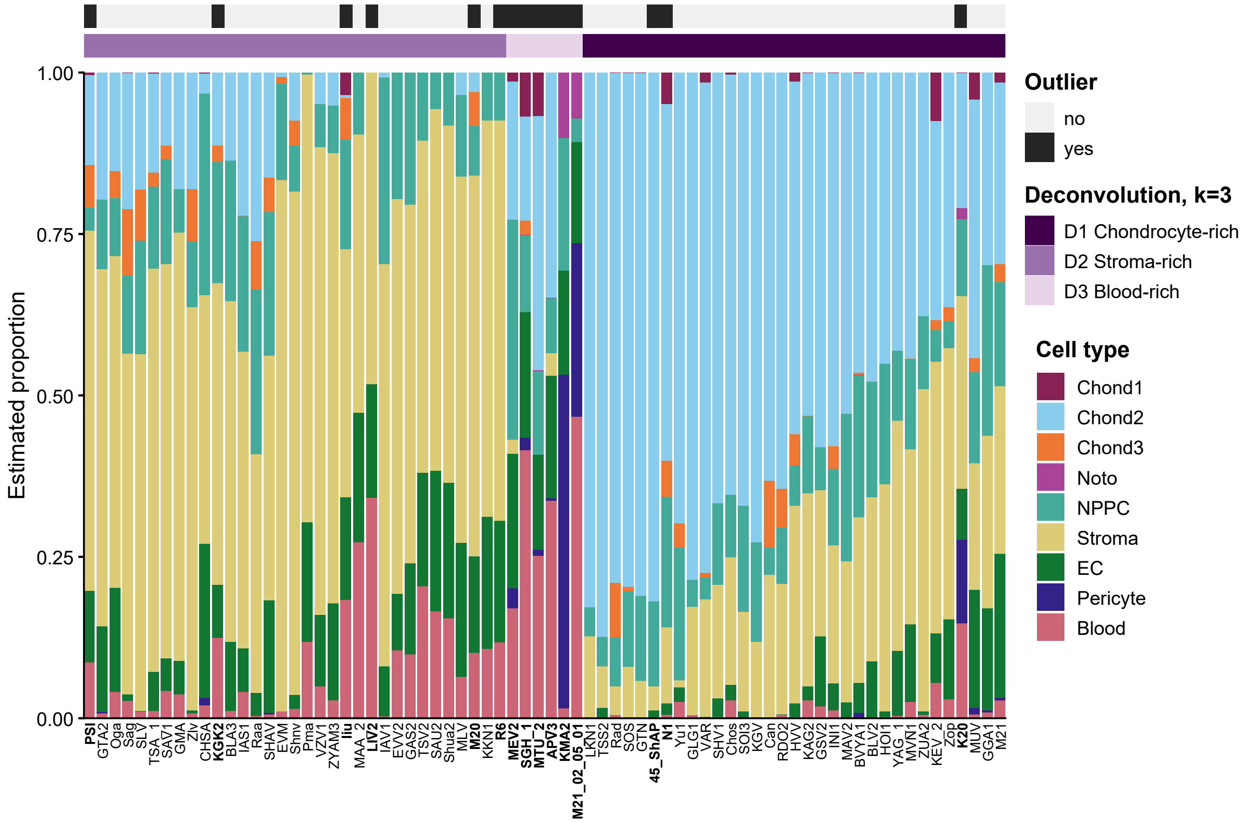


**Supplementary Fig. 2 | Cell-type deconvolution of bulk RNA-seq samples.** Stacked bars show the BayesPrism-estimated proportions of the nine reference cell types in each of the 72 samples. Samples are ordered by the hierarchical clustering of their cell-type proportion vectors, cut at $k = 3$ into three deconvolution groups named by composition: D1, Chondrocyte-rich (highest mean Chond2 proportion); D2, Stroma-rich (highest mean Stroma proportion); and D3, Blood-rich. The upper strip marks the 15 outlier samples excluded at quality control, whose labels are also shown in bold; the lower strip shows the deconvolution group. Noto: notochordal cells; NPPC: nucleus pulposus progenitor cells; Stroma: stromal cells; EC: endothelial cells.

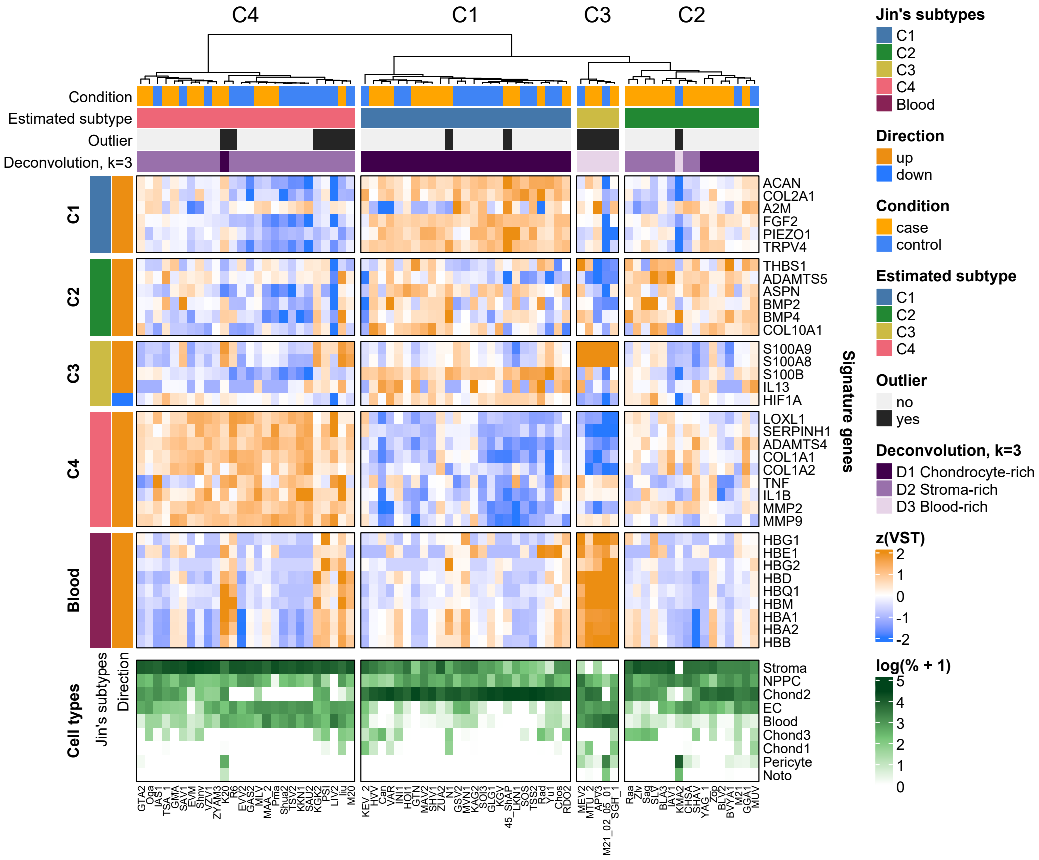


**Supplementary Fig. 3 | Expression of Jin et al. subtype signature genes and haemoglobin genes across all 72 RuDDS samples.** Samples were clustered using Pearson correlation distance computed over the Jin et al. classifier genes (Supplementary Fig. 4), and the dendrogram was cut at $k = 4$. Each cluster is labelled with the Jin subtype whose marker and classifier genes it expresses most highly: C4 ($n = 26$), C1 ($n = 25$), C3 ($n = 5$) and C2 ($n = 16$). The upper heatmap shows per-gene z-scores of variance-stabilised expression for subtype signature genes and for nine haemoglobin genes (Blood block). The signature genes are the representative subtype-specific differentially expressed matrisome genes of Jin et al. (their Fig. 2e), except *COL9A1*. The Direction bar shows whether each marker gene is expected to be up- or downregulated in its subtype in the reference study. The lower heatmap shows the BayesPrism cell-type proportions of the same samples as log(% + 1). Column annotations, from top to bottom: Condition, case or control; Estimated subtype; Outlier, samples excluded at quality control; Deconvolution, $k = 3$, the deconvolution groups of Supplementary Fig. 2. The haemoglobin genes were not used for clustering. All five samples of the C3 cluster are outliers from the Blood-rich deconvolution group and show coordinated high expression of the haemoglobin genes together with *S100A8* and *S100A9*, which are C3 markers in Jin et al. but are also abundant in blood neutrophils and monocytes.

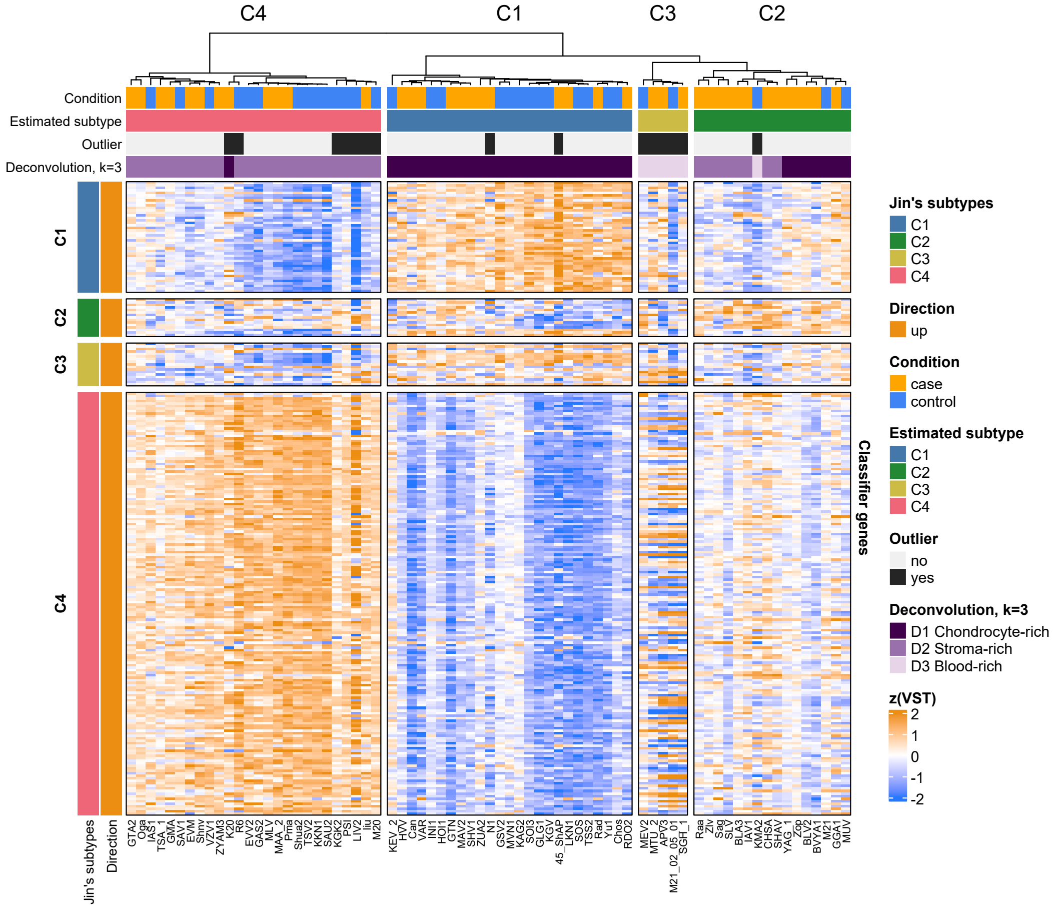


**Supplementary Fig. 4 | Expression of the Jin et al. classifier genes across all 72 RuDDS samples.** The heatmap shows per-gene z-scores of variance-stabilised expression (DESeq2 VST) for the 244 genes of the 248-gene Jin et al. subtype classifier panel that are present in the RuDDS data; this matrix was used to cluster the samples in Supplementary Fig. 3. Rows are grouped by the subtype to which Jin et al. assigned each gene as upregulated. Columns follow the dendrogram, $k = 4$ split and annotations of Supplementary Fig. 3.

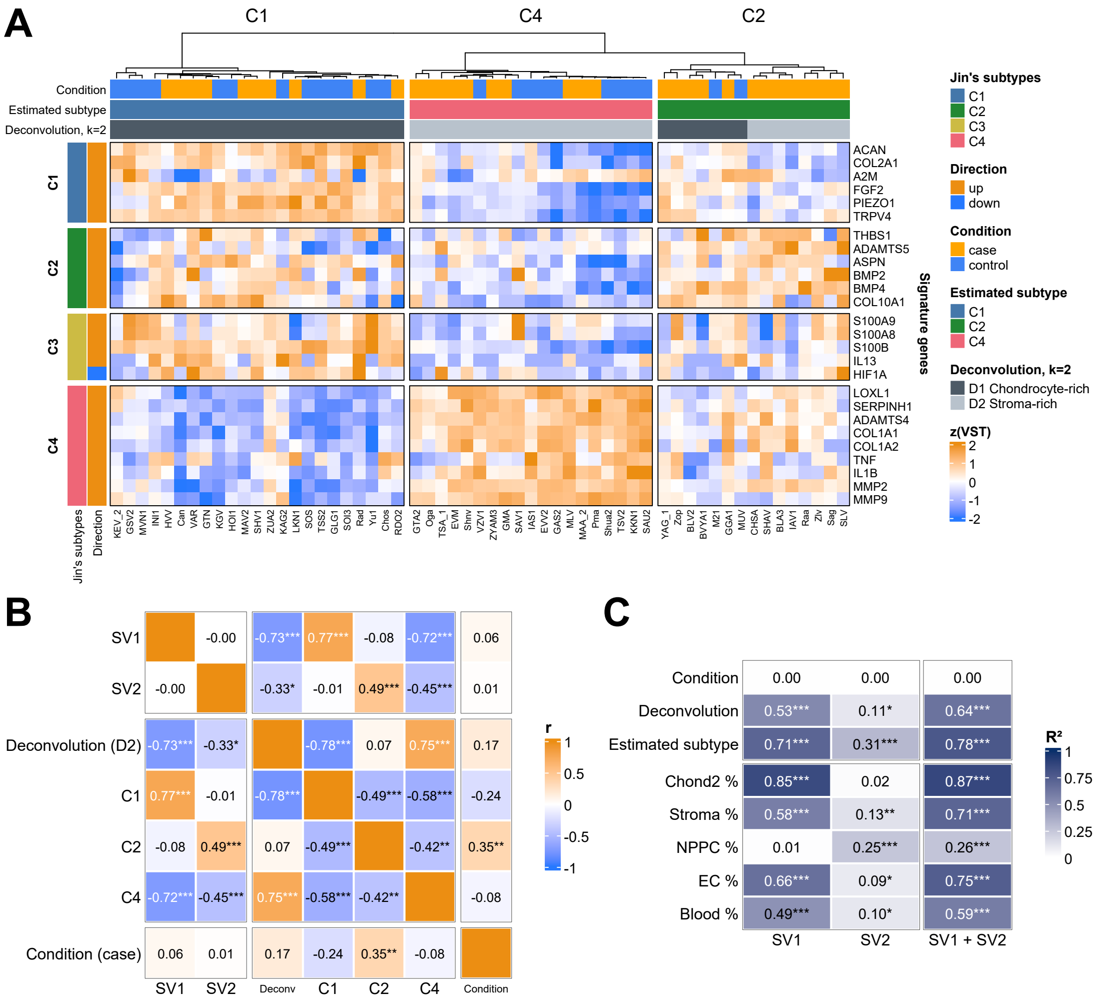


**Supplementary Fig. 5 | Estimated subtypes and deconvolution groups of the 57 samples passing quality control, and their association with the surrogate variables of the DESeq2 model.** (A) Expression of the subtype signature genes, shown as in Supplementary Fig. 3. Clustering on the classifier genes was repeated in the 57 samples and cut at $k$ = 3: C1 ($n$ = 23), C4 ($n$ = 19) and C2 ($n$ = 15). Deconvolution, $k$ = 2: clustering of the cell-type proportions of the 57 samples into D1, Chondrocyte-rich, and D2, Stroma-rich groups. (B) Pearson correlations between the surrogate variables (SV1, SV2), deconvolution group (D2), estimated subtype (C1, C2, C4) and condition. Each binary variable is coded 1 for D2, for the given subtype or for cases, and 0 otherwise. The subtype variables are mutually exclusive and therefore correlate negatively by construction. Because SV signs are arbitrary, only the magnitude of correlations with an SV is interpretable. (C) Variance shared ($R^2$) between each variable and SV1, SV2, or both SVs jointly. For a single SV, $R^2$ is that of a linear regression of the SV on the variable ($\eta^2$ for groupings). For SV1 + SV2, it is the $R^2$ of a regression of the variable on both SVs (the squared first canonical correlation for estimated subtype). The upper block shows the groupings and the lower block the estimated proportions (%) of five cell types. Nominal $P$-values: correlation test (B); $F$ test for a single SV and Pillai's trace MANOVA for SV1 + SV2 (C). $^{*}P < 0.05$, $^{**}P < 0.01$, $^{***}P < 0.001$. Jointly, SV1 and SV2 account for 64% of the variance in deconvolution group and 87% and 71% in Chond2 and Stroma proportions.

**Supplementary Note 1. Sample outlier detection.**

Four samples (45_ShAP, K20, N1, R6) were sequenced under an earlier pilot library-preparation protocol. Compared with the remaining 68 samples (two-sided Wilcoxon rank-sum test), these samples had a lower proportion of reads not overlapping any feature (4.2% vs. 9.0%, $P$ = 0.002) and fewer detected genes ($\sim$19,800 vs. $\sim$21,600, $P$ = 0.004). These metrics are concordant with their separation on PC3 and PC4 (Supplementary Fig. 1B). Sample PSI, recorded as male, expressed Y-linked genes together with the female-specific transcript *XIST* at a level typical of female samples, indicating probable contamination with RNA from a female donor.

LIV2 and KMA2 differed substantially from the rest of the cohort (Supplementary Fig. 1B, C). KMA2 was excluded on the basis of PC1/PC2 and PC5/PC6 clustering (Supplementary Fig. 1A, C) and a pericyte proportion far above that of the other samples (0.52 vs. a mean of 0.007; Supplementary Fig. 2). LIV2 was excluded on the basis of PC1/PC2, PC3/PC4 and PC5/PC6 clustering (Supplementary Fig. 1A–C).

The remaining eight outliers (APV3, liu, KGK2, M20, M21_02_05_01, MEV2, MTU_2, SGH_1) were excluded on the basis of PC1/PC2 clustering (Supplementary Fig. 1A) and also showed elevated expression of haemoglobin genes (Supplementary Fig. 3).

**Supplementary Note 2. Count outlier detection using Cook's distance.**

In RNA-seq data some genes may show unexpectedly high counts in a single sample. To address this issue, DESeq2 calculates Cook's distance for each gene and for each sample, showing the impact of an individual sample on the fitted coefficients for this gene. If the design formula contains only categorical variables, DESeq2 automatically determines the optimal Cook's distance threshold, above which an expression value would be considered an outlier. Given that our initial design formula included one continuous variable (Age), we calculated Cook's distance threshold manually to identify only the most extreme count outliers to avoid overcorrection. A gene count value was flagged as an outlier if its Cook's distance exceeded the upper quartile of that gene's Cook's distances by more than 50 interquartile ranges. Such extreme outliers among the count values (0.02% of all values) were assigned the weight of 0, using DESeq2's capability to downweight individual data points.
